# Coinfection of semi-infectious particles can contribute substantially to influenza infection dynamics

**DOI:** 10.1101/547349

**Authors:** Alex Farrell, Christopher Brooke, Katia Koelle, Ruian Ke

## Abstract

**Abstract:** Influenza is an RNA virus with a genome comprised of eight gene segments. Recent experiments show that the vast majority of virions fail to express one or more gene segments and thus cannot cause a productive infection on their own. These particles, called semi-infectious particles (SIPs), can induce virion production through complementation when multiple SIPs are present in an infected cell. Previous within-host influenza models ignore the potential effects of coinfection and SIPs during virus infection. Here, to investigate the extent that SIPs and coinfection impact viral dynamics, we constructed two within-host models that explicitly keep track of SIPs and coinfection, and fitted the models to clinical data published previously. We found that the model making a more realistic assumption that viruses can only reach a limited number of target cells allows for frequent co-infection during early viral exponential growth and predicts that SIPs contribute substantially to viral load. Furthermore, the model provides a new interpretation of the determinants of viral growth and predicts that the virus within-host growth rate (a measure of viral fitness) is relatively insensitive to the fraction of virions being SIPs, consistent with biological observations. Our results highlight the important role that cellular co-infection can play in regulating infection dynamics and provide a potential explanation for why SIP production is not highly deleterious. More broadly, the model can be used as a general framework to understand coinfection/superinfection in other viral infections.

**Author Summary:** Influenza A viruses (IAVs) represent a large public health burden across the world. Currently, our understanding of their infection dynamics is incomplete, which hinders the development of effective vaccines and treatment strategies. Recently, it was shown that a large fraction of virions, called semi-infectious particles, do not cause productive infection on their own; however, coinfection of these particles leads to productive infection. The extent that semi-infectious particles and, more broadly, coinfection contribute to overall influenza infection dynamics is not clear. To address this question, we constructed mathematical models explicitly keeping track of semi-infectious particles and coinfection. We show that coinfection can be frequent over the course of infection and that SIPs play an important role in regulating infection dynamics. Our results have implications towards developing effective therapeutics.

## Introduction

Influenza A viruses (IAVs) cause hundreds of thousands of hospitalizations, tens of thousands of deaths, and cost tens of billions of dollars each year in the United States alone [1]. In addition, pandemic strains emerge from time to time from reassortment of human, swine and/or bird strains, leading to high morbidity and mortality rates [2–4]. Extensive efforts have been made to understand the transmission and evolution of IAVs at the epidemiological level [5,6]; however, our understanding of the molecular origins of within-host genomic diversity of IAVs, and how this diversity affects infection dynamics is incomplete [7]. This knowledge may be crucial for understanding within-host IAV evolution, the frequency of *in vivo* reassortment, and ultimately for the development of effective vaccines and treatment strategies [7,8].

IAVs are negative sense RNA viruses with genomes comprised of eight gene segments. Proteins from all eight segments are essential for completion of the viral replication cycle. Recently, Brooke and colleagues reported that the majority of virions (between 70% and 98%) fail to express a complete set of genome segments, and thus are non-infectious under traditional limiting-dilution assays [9]. These non-infectious virions are broadly categorized as ‘semi-infectious particles’ (SIPs), whereas in contrast, virions expressing all eight segments are termed ‘fully infectious particles’ (FIPs). If two SIPs infect the same cell, it is likely that they will complement each other to express at least one copy of all eight gene segments and thereby be capable of producing viral progeny that will go on to infect other cells [8,10]. This phenomenon, called ‘multiplicity reactivation’ or ‘complementation’ provides SIPs the potential to induce a productive infection within a cell in the absence of FIPs. Importantly, it was shown that the presence of SIPs enhances IAV reassortment and diversity and thus may accelerate the rate of IAV evolution [11]. However, the extent to which coinfection of SIPs occurs and contributes to the overall viral dynamics is not clear.

Previously, mathematical models have been used to describe IAV viral load dynamics [12], estimate drug efficacies [13,14], and characterize the role of innate and adaptive immunity [15–17]. While some models have included ‘non-infectious’ virions [18–20], the roles that coinfection and complementation between SIPs play in regulating within-host IAV dynamics has not been studied. This knowledge is crucial to a quantitative understanding the extent of co-infection and reassortment during infection [11]. To address this gap, we constructed two viral dynamic models that differ in their assumptions of the infection process. We identified the conditions that allow SIPs to be ‘biologically active’, i.e. inducing productive infection, and examined the role of complementation between SIPs in driving within-host viral growth. We show that ignoring SIPs may lead to imprecise estimates of parameter values (when fitting models to data) and an incomplete understanding of the factors that drive viral load dynamics. In addition, we show that a model considering frequent co-infection and SIPs can reproduce key features of existing experimental and clinical data. We find that in this model, the exponential viral growth rate is insensitive to the relative ratio of FIP to SIP production, suggesting that the production of large numbers of SIPs may not compromise overall replicative capacity. In general, our model may also be useful to understand the roles and consequences of coinfection/superinfection in other viral infections [21].

## Methods

The homogenous mixing (HM) model. We first constructed a model including SIPs and coinfection dynamics assuming viral particles and cells are homogenously mixed. This model is an extension of a basic within-host viral infection model that assumes target-cell limitation [12]. It is described by the following system of ordinary differential equations (ODEs):

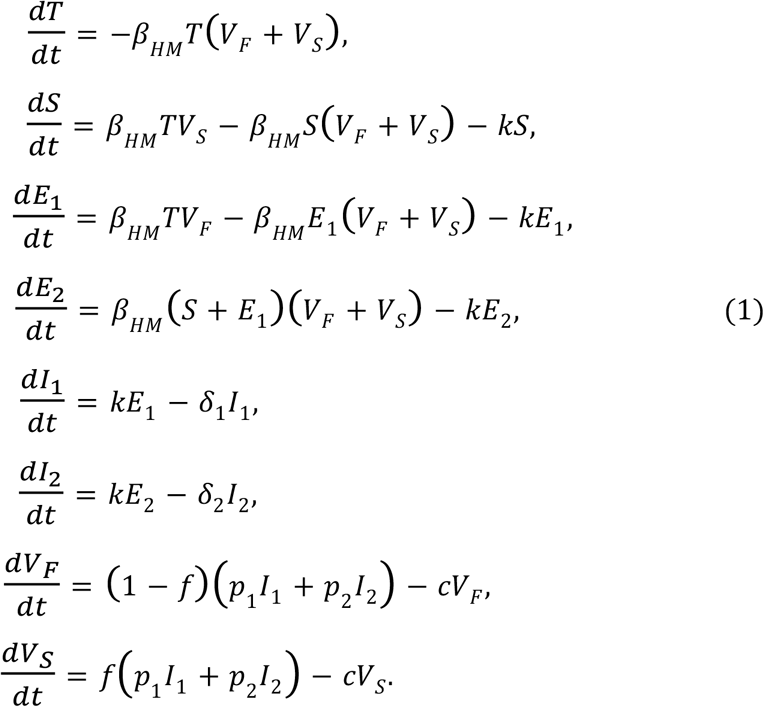

In this model, target cells (*T*) are infected with FIPs (*V_F_*) and SIPs (*V_S_*) to become, respectively, FIP infected cells (*E_1_*) and SIP infected cells (*S*) at rate *β_HM_*. Note that viral infection events are modeled using a mass-action term in this model, i.e. viruses and cells are well-mixed. This assumption implies that once a virion is produced, it has equal probability to contact every cell in a host. Although this assumption is often used in models for simplicity [12,16,22], it is clearly biologically unrealistic, especially for influenza infection where target cells are distributed spatially from upper to lower respiratory tract [23].

FIP infected cells go through an eclipse phase (*E_1_* cells) during which no virions are produced. *E_1_* cells then mature to singly infected virion producing cells, *I_1_*, at rate *k*. It has been shown that a cell co-infected with two SIPs will likely induce viral production in the cell through complementation, i.e. multiplicity reactivation [8]. Thus, we assume that if *E_1_* or *S* cells are coinfected by either a fully or semi-infectious virion, they become doubly infected cells in an eclipse phase, *E_2_*, which then mature to virion producing cells, *I_2_*, at rate *k*. For SIP infected cells (*S*), there exists a period of time (1/*k* on average) when they can become superinfected. After this period, the cells become resistant to superinfection, and thus these cells are removed from our system [24,25]. Singly and doubly infected virion producing cells (*I_1_* and *I_2_*, respectively) die at per capita rate *δ_1_* and *δ_2_*, respectively. We do not consider superinfection of *I_1_* or *I_2_* cells, because it has been shown that only cells in their eclipse phase are vulnerable to superinfection [24]. Virions are produced from singly and doubly infected virion producing cells at rates *p_1_* and *p_2_* respectively. We expect that the viral output from doubly infected cells is higher than that from singly infected cells, i.e. *p_2_* > *p_1_* [26,27]. Of all virions produced, a constant fraction [9], *f*, of them are semi-infectious, while the remaining fraction are fully infectious. Both FIPs (*V_F_*) and SIPs (*V_S_*) are cleared at rate *c*.

### The target cell saturation (TS) model

We considered an alternative model using Michaelis-Menten (also known as Holling Type II) functions to model infection of susceptible cells. This type of infection term has been used previously in HIV within-host models [28,29] and can arise in situations where cells are not well mixed, for example, when target cells are spatially segregated. Here, our interpretation of the Michaelis-Menten term is that during infection a virus cannot reach all target cells in a host; instead, there are only a small number of neighboring target cells that a virus can physically access. Thus, the infection ‘saturates’ i.e. there is a maximal infectivity possible that is effectively independent of the number of susceptible cells so long as the number of susceptible cells is large, which is described by the Michaelis-Menten term. For virus infection of uninfected cells (*T*) we use the term 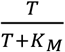, where *K_M_*, the saturation parameter, describes the amount of cells at which half of the maximum infectivity is reached. In this case, there are a limited number of target cells that can be infected (defined by *K_M_*); when the number of susceptible cells is greater than *K_M_*, the rate of infection of susceptible cells saturates to its maximum. For infection of already infected cells, i.e. *S, E_1_*, and *E_2_*, the denominator of the Michaelis-Menten term is set to 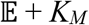, where 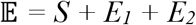. This is motivated by the idea that already infected cells are close to each other at an infection site (especially when the infection process is spatially structured [30]), so a virion that is produced from productively infected cells can reach those eclipse phase cells in the neighborhood. Thus, we sum up all the eclipse phase cells that can be superinfected. The system of ODEs for the TS model is shown below. The only differences between this model and the HM model are the Michaelis-Menten infection terms.

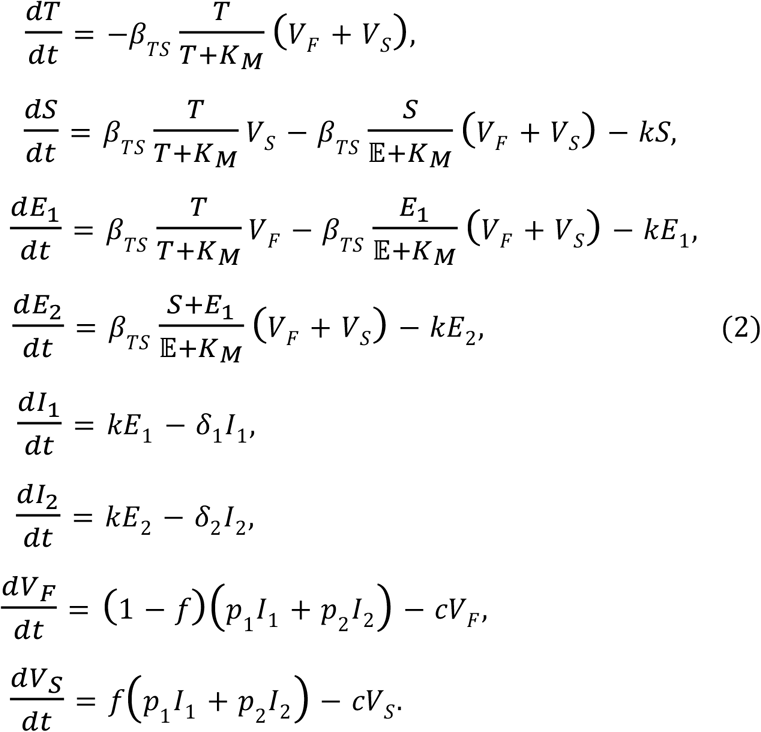

The assumption of target cell saturation is more realistic, especially for influenza infections in tissue. For example, it has been shown in human tracheobronchial epithelium that foci of infected cells form during infection and a virion produced may be more likely to infect cells that are closer to the cell that produced the virion [30]. Similar results have also been observed in the lungs of mice [31–33] and ferrets [34]. Since virions are produced within such a focus of infection, their neighboring cells are more likely to have already been infected, leading to rapid co-infection.

Note that because of the Michaelis-Menten terms we use, the units and interpretation of the parameter *β_TS_* differ from *β_HM_*. Here, *β_TS_* is the maximum infectivity rate per virion when target cells reach saturation. A schematic of both models is shown in Fig. 1.

**Fig. 1.**
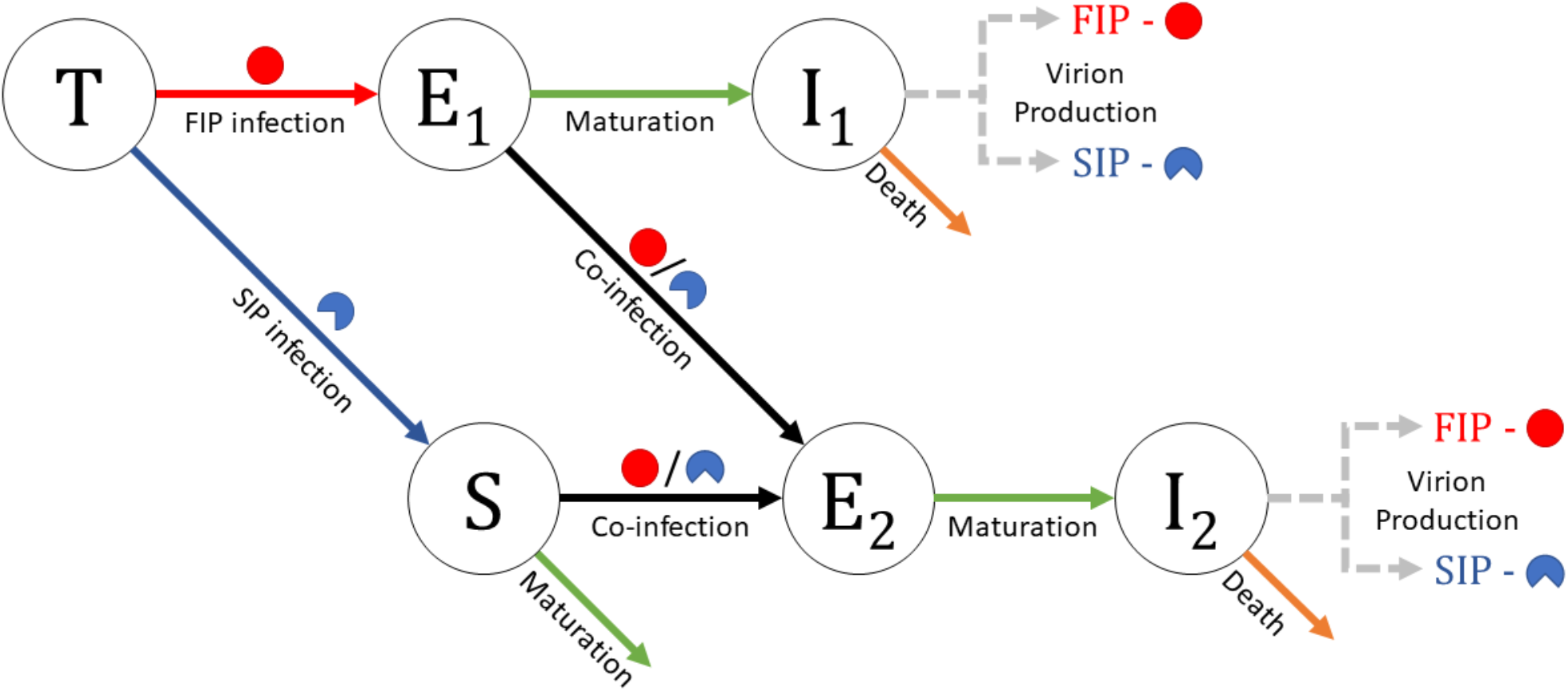
A schematic of the within-host influenza models that include semi-infectious particles (SIPs). Fully infectious particles (FIPs) are represented by red circles and SIPs by blue partial-circles. Uninfected target cells (*T*) can be infected by FIPs and SIPs to become cells in an eclipse phase (*E*_1_ and *S*, respectively). When cells in an eclipse phase (*S* or *E*_1_) are further infected by FIPs or SIPs, they become co-infected cells in an eclipse phase (*E*_2_). *E*_1_ and *E*_2_ cells mature into productively infected cells (*I*_1_ and *I*_2_, respectively) to produce FIPs and SIPs.

### Parameter values and data fitting procedure

To determine the abilities of the HM and TS models to capture observed patterns of viral load, we fit both models to viral load data from a human challenge study that was described in [35] and first analyzed in [12]. The fitting procedure was performed using a built-in, derivative free simplex search method in MATLAB. We set *p_2_* = 2_*p1*_ when fitting our model to the data. We further fixed several parameters. We set the initial target cell count to *T*(0)=4×10^8^ cells, based on [12], and the maturation time *1/*k** to 1/8 days (3 hours), based on Ref. [24]. We further set c = 15 d^-1^, and *f* = 0.9 (dimensionless) as biologically reasonable parameter values for the virion clearance rate [36] and fraction of virions produced that are semi-infectious [8,9,11], respectively. The remaining parameters of infectivity rate, saturation parameter (for TS model), death rates of virion producing cells, virion production rate and initial FIP concentration, i.e. *β_HM_* (or *β_TS_* and *K_M_*, depending on the model), *δ_1_, δ_2_, p_1_,* and V_F_(0) are fit to the data. We calculated the initial SIP concentration, *V_S_*(0), by assuming that the viral input dose maintained the constant proportion of FIPs to SIPs seen in viral production, i.e. 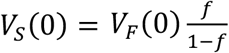 We note that TCID_50_ viral titers are measurements of fully infectious virus only, thus we fit *V_F_* to the data. While fitting the model to data, we restricted *p_1_* to be less than or equal to 10 TCID50/ml× d^-1^, *K_M_* to be greater than or equal to 1 cell and bounded *V_F_*(0) above and below by 1 and 0.001 TCID50/ml respectively. Finally, we assume that δ1 ≤ δ2.

## Results

### Fitting the models to published *in vivo* data

We constructed two mathematical models incorporating SIPs, i.e. the homogenous mixing (HM) model and the target cell saturation (TS) model (see Methods). The HM model follows previous model assumptions that the virus and cells are well mixed; whereas the TS model assumes that there is only a small number of target cells that a virus can reach. We first fitted the two models to viral load data reported in Ref. [12] and collected in Ref. [35], where six serosusceptible adults were voluntarily infected with a cloned influenza A virus, followed by daily nasal washes for one week to measure the patients’ viral titers. Both models describe the data set very well (see Table 1 for the best fit parameters for each patient in Fig. 2). They captured the exponential viral growth during the first 2-4 days of infection and the exponential decline after viral peak. In patient 2 and 5, both models predicted two phases of viral decline as suggested by the data (see below for details). Although both models describe the clinical data, below we show that these two models lead to very different predictions about the infection dynamics, and we found that the TS model is a better model to recapitulate a wide range of other biological evidence.

**Table 1.** Best fit parameters for both models to data in Ref. [12]: HM is the best fit for the homogeneous mixing model and TS is for the target saturation model. Res. is the residue (i.e. sum of square error). The HM model does not contain the parameter *K_M_*. The units of β are (TCΓD50/ml)^-1^d^-1^cell^-1^ for the HM model and (TCID50/ml)^-1^d^-1^ for the TS model. The units of *p*_1_ are TCID_50_/ml× d^-1^. See Methods for further details on the fitting procedure.

| Patient | Model | $\beta$ | $K_M$ (cells) | $\delta_1$ ($\text{d}^{-1}$) | $\delta_2$ ($\text{d}^{-1}$) | $p_1$ | $V_F(0)$<br>($\text{TCID}_{50}/\text{ml}$) | Res. |
| --- | --- | --- | --- | --- | --- | --- | --- | --- |
| 1 | HM | $9.55 \times 10^{-6}$ | N/A | 2.85 | 3.22 | 0.975 | 0.435 | 4.9 |
|  | TS | 895 | 1.04 | 3.03 | 3.03 | 0.548 | 0.142 | 4.8 |
| 2 | HM | $6.93 \times 10^{-5}$ | N/A | 1.59 | 7.18 | 0.265 | 0.001 | 4.3 |
| | TS | $5.03 \times 10^3$ | 1 | 1.55 | 8.33 | 0.229 | 0.001 | 4.3 |
| 3 | HM | $1.63 \times 10^{-5}$ | N/A | 1.98 | 2.16 | 0.436 | 1 | 8.9 |
| | TS | $1.16 \times 10^3$ | 1.02 | 2.1 | 2.12 | 0.289 | 1 | 8.5 |
| 4 | HM | $1.01 \times 10^{-6}$ | N/A | 3 | 3.04 | 10 | 1 | 4.2 |
|  | TS | 57.6 | 3.71 | 3.08 | 3.08 | 7.9 | 1 | 3.8 |
| 5 | HM | $7.15 \times 10^{-7}$ | N/A | 4.27 | 7.06 | 10 | 1 | 3.3 |
|  | TS | 43.6 | 1 | 4.72 | 9.08 | 9.95 | 1 | 3.0 |
| 6 | HM | $6.45 \times 10^{-7}$ | N/A | 3.67 | 4.55 | 10 | 1 | 14 |
|  | TS | 32.1 | 1 | 5.23 | 4.47 | 10 | 1 | 10.9 |

**Fig. 2.**
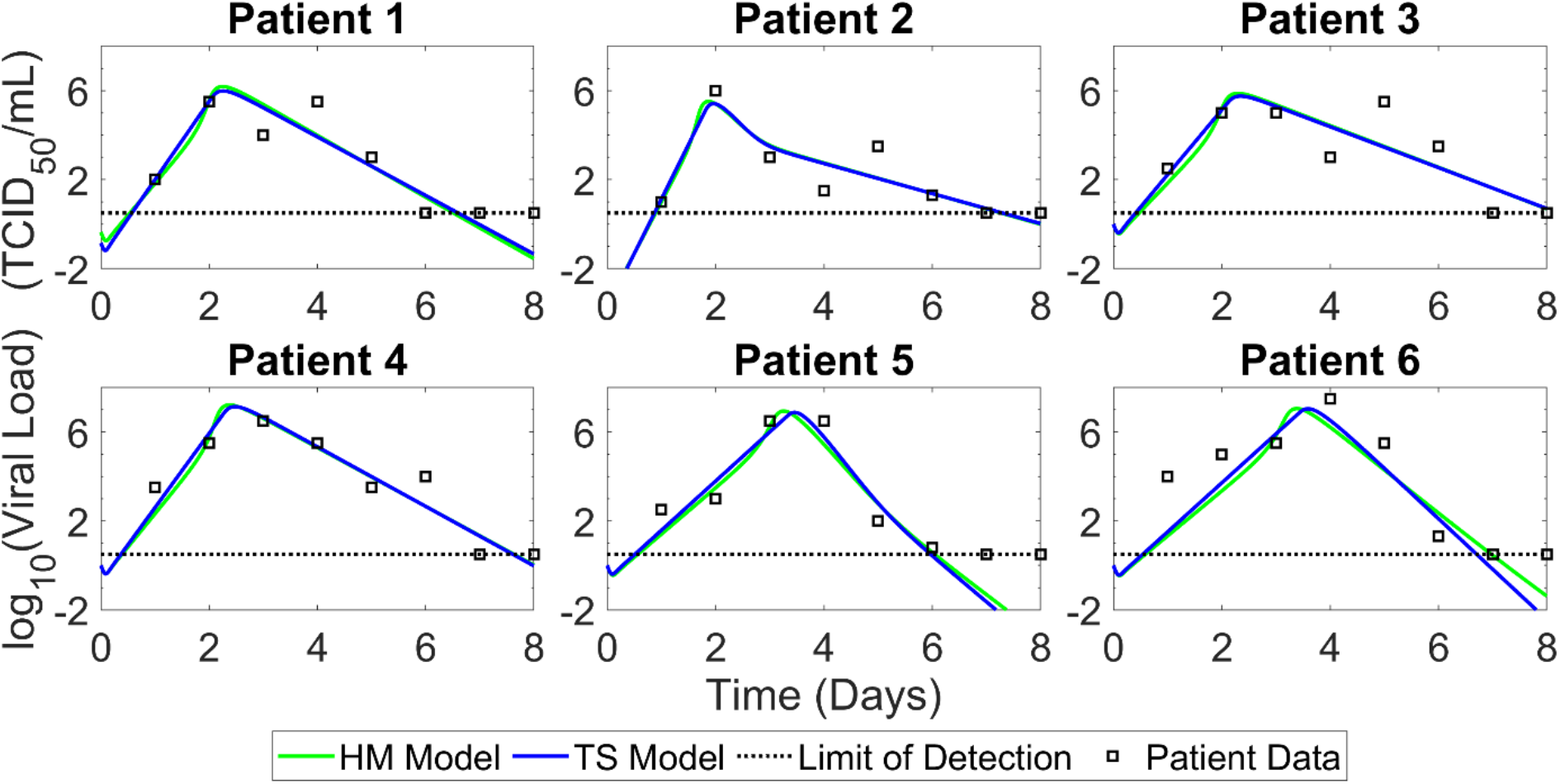
Fitting results of both the HM model and the TS model to clinical data (from Ref. [12]). Results are shown for data from 6 participants. Best fits of the HM model and the TS model are shown as blue and green lines, respectively. Data points are shown as squares and the dotted line is the limit of detection of the experiment. The parameters for each patient are presented in Table 1.

### Viral load dynamics and frequency of co-infection

We first analyzed the infection dynamics predicted by the HM model. More specifically, we kept track of the FIPs produced from singly infected cells (I_1_) and coinfected cells (I_2_), and the frequency of co-infected cells (I_2_+E_2_) among all infected cells (S+E_1_+I_1_+E_2_+I_2_). In addition, to investigate the extent of the contribution of ‘multiplicity reactivation’ (due to complementation of SIPs) towards the viral load, we extended the models to keep track of FIPs produced as a result of ‘multiplicity reactivation’ (see Fig. S1 for a schematic for the extended model).

The HM model predicts that much of the initial infection period is driven by the exponential growth of viruses produced from singly infected cells (Fig. 3A and B), similar as previous models [12,16,22]. This is because the number of uninfected cells is orders of magnitude higher than other cell classes, and a virus will mostly likely encounter an uninfected cell under the homogenous mixing assumption. Then, there exists a brief period before the viral peak where the number of already infected cells exceeds the number of uninfected cells, co-infection frequency rapidly increases, and doubly infected cells begin to meaningfully contribute to the viral load. Therefore, the HM model predicts an extremely low frequency of co-infection during most of the exponential growth period (Fig. 3B), which is inconsistent with experimental data [8,37,38]. Further, this prediction indicates that SIPs do not significantly contribute to exponential viral growth period until peak viremia.

**Fig. 3.**
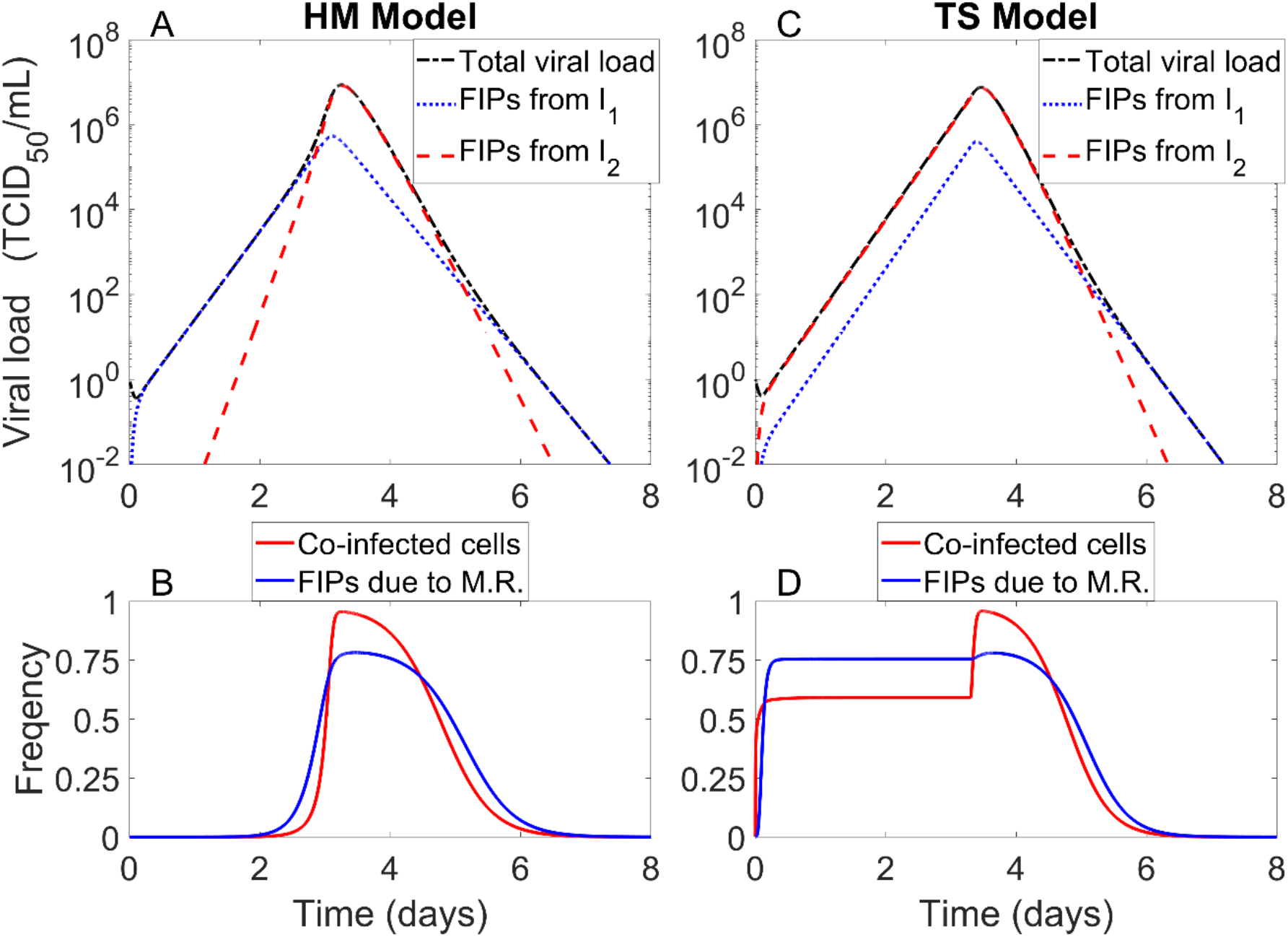
Distinct predictions about the drivers of viral load dynamics and the frequency of co-infected cells by the HM model and the TS model. **(A)** Viral load dynamics and drivers of them predicted by the HM model. The total viral load is the concentration of fully infectious virus (FIPs) from all sources. FIPs from *I*_1_ and *I*_2_ (blue and red lines, respectively) are the densities of fully infectious virus produced by singly and doubly infected virion producing cells, respectively. **(B)** The frequency of infected cells that are co-infected (the red line) and the frequency of fully infectious virions produced that are due to multiplicity reactivation of SIPs (the blue line; calculations for this quantity are explained in caption of Fig. S2). Multiplicity reactivation occurs when a SIP co-infects a cell already infected by a SIP (*S*). The parameters used are from patient 5 in Table 1. (**C and D**) Predictions of the viral load dynamics of frequency of coinfection from the TS model. The TS model predicts the viral load is primarily driven by FIP production from coinfected cells and the frequency of coinfection remains high throughout most of the infection course.

In contrast to the single exponential growth phase as predicted by the HM model, the TS model can predict two phases of exponential growth (Fig. S2A). The first phase is driven by FIP production from singly infected cells. During this phase, the viral load is low, the levels of the *E*_1_ and *S* cells (targets for co-infection) are very low and thus coinfection is infrequent (Fig. S2B). When the viral load increases to a level such that the levels of the E1 and the *S* cells increased, coinfection becomes frequent and the viral load is driven by viral production from co-infected cells, i.e. the 2^nd^ second phase of exponential growth. During this phase, both FIPs and SIPs contribute to viral growth. The transition from the 1^st^ to the 2^nd^ growth phase is determined by the value of the saturation parameter, *K_M_*, relating to the total number of target cells that a virus can reach (Fig. S3). Fitting the model to clinical data showed that *K_M_* is very small (Table 1), suggesting that the number of target cells that a virus can reach is very low compared to the total number of target cells in a human body, This is a reasonable estimate because influenza infections occur in tissue, where cells are spatially segregated and it is likely that the number of cells that a virus can contact is very limited. In this case, the TS model predicts that the 1^st^ phase growth lasts for a very short period of time and consequently co-infection occurs frequently during most of infection period (Fig. 3C and D), consistent with previous experimental studies [37–40].

In both the HM and TS models, the viral decline after peak viremia is due to the depletion of target cells. When the doubly infected cells die at a higher rate than the singly infected cells, we would expect a bi-phasic viral decline, consistent with findings of our recent study [41]. This bi-phasic viral decline is consistent with data sets from human challenge studies and experimental studies on ponies [12,15]. Our model simulations (Fig. 3A and C) show that at peak viremia most infected cells are doubly infected, and thus the first phase of decline is driven by death of the doubly infected cells. Over time, doubly infected cells are cleared, singly infected cells become dominant and the viral decline is then driven by singly infected cells.

Overall, we argue that the TS model makes more realistic assumptions about the infection process of influenza virus in a host compared to the HM model, i.e. viruses and cells are not well-mixed and factors, such as spatial segregation, limit the number of target cells that a virus can reach. With the more realistic assumption, the TS model is able to predict co-infection frequencies that are consistent with biological observations [8,37,38].

We then further quantified the contribution of SIPs to viral load through ‘multiplicity reactivation’ or ‘complementation’, i.e. productive infection by two SIPs [8]. Using the extended HM and the TS model (the schematic is shown in Fig S1), we kept track of cells that are infected with two SIPs. We found that the TS model predicts that SIPs alone can contribute substantially to the fully infectious viral load while the viral load is growing, contributing nearly 75% of all FIPs produced during that time (Fig. 3D), in stark contrast to the predictions of the HM model (Fig. 3B).

### Dependence of viral growth rate on the proportion of SIPs produced

Depending on the IAV strain, SIPs can compose between 70% and 98% of biologically active particles in an IAV population [9]. While the ratio of virions produced that are semi-infectious is not the only difference between strains, the large variation in the fraction of SIP production suggests that this fraction is not an important determinant of viral fitness [9,39]. One measure of *in vivo* viral fitness is the rate of exponential growth. We thus simulated both the HM model and the TS model using realistic parameter values and examined how strongly the viral growth rate (at day 1 post-infection) depends on the fraction of SIP production. Simulation results show dramatic differences between the two models’ predictions (Fig. 4).

**Fig. 4.**
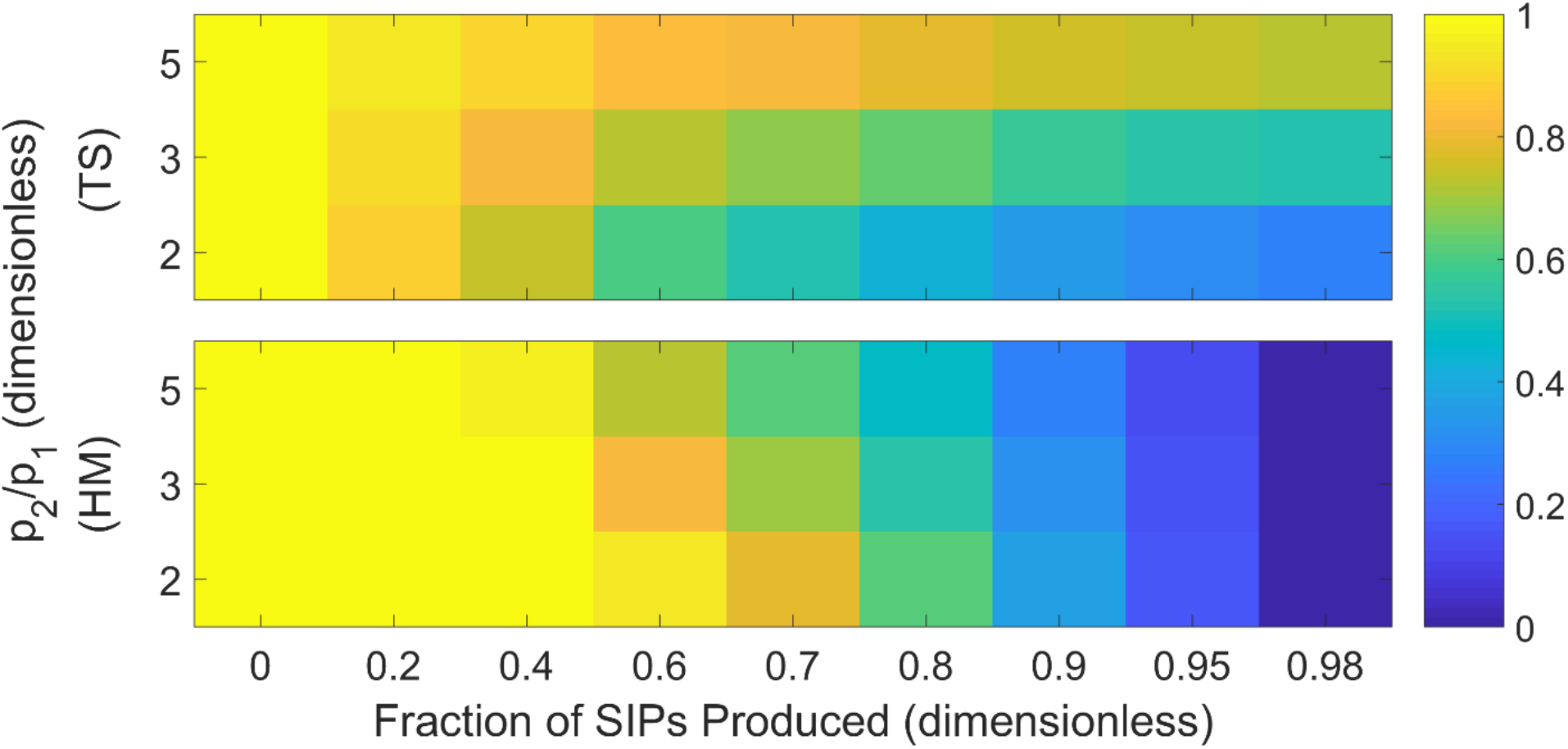
Sensitivity/insensitivity of the exponential viral growth rate to the fraction of SIPs produced from infected cells depends on the model structure. Simulated viral growth rates, measured at one day post infection, are compared over a range of values for the fraction of virions produced that are semi-infectious (x-axis). We chose multiple values for the rate of virion production from cells that are doubly infected (2, 3 and 5), keeping the virion production from singly infected cells constant. For a fair comparison, we normalize all growth rates by their value when all virions produced are fully infectious. The parameters used in these calculations are from patient 5 in Table 1, except for *p*_2_.

The HM model predicts that the viral growth rate is driven by FIPs produced from singly infected cells. The rate of exponential growth in the HM model, denoted by *λ_HM_*, can be approximated by (see S1 Supporting Information):

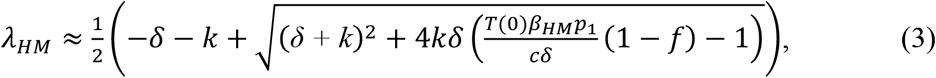

where *T*(0) is the equilibrium concentration of target cells in the absence of infection. This expression shows that the rate of exponential growth, *λ_HM_*, is extremely sensitive to changes in the fraction of SIP production, *f* The higher fraction of SIP production, the lower FIP production and thus the lower the growth rate.

In contrast, the TS model predicts that the growth rate is relatively insensitive to changes in the fraction of SIP production, especially when co-infected cells produce more virions than singly infected cells (i.e. higher ratios of *p*_2_/*p*_1_ in Fig.4). This is because with a large fraction of SIP production, SIP can contribute to the viral growth through complementation. We analytically approximated the rates of the two exponential growth phases (see S1 Supporting Information):

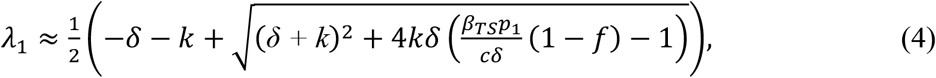

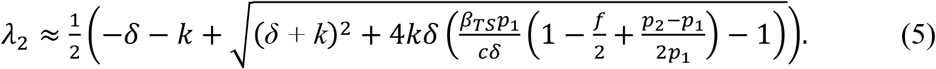

Eqn. 5 shows that the exponential rate of the second phase viral growth (which dominates the exponential growth period based on our model fitting), *λ*_2_, is less sensitive to changes in *f*, especially when 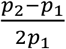 is large, i.e. the total number of viral particles produced from coinfected cells is larger than the number of particles produced from singly infected cells.

These results again argue that the TS model is more plausible model than the HM model and other simpler models that do not include co-infection [12,16,22]. The TS model provides a new description of the exponential growth observed in viral load data (Eqns. 4 and 5). In addition, Eqn. 5 also suggests that the viral growth rate in the TS model is dependent on viral production rates from both singly and co-infected cells, highlighting the importance of quantifying these parameters experimentally.

## Discussion

In this study, we constructed mathematical models explicitly considering co-infection dynamics and the effects of SIP production during influenza virus infection. By comparing two alternative models against biological evidence, we demonstrated the importance of incorporating realistic assumptions about the infection process. Using the more realistic model that assumes that a virus can only reach a limited number of target cells, we showed that coinfection is frequent during influenza infection in a host and consequently SIPs can contribute substantially to and regulate viral load dynamics through complementation. Our model is a useful tool to understand coinfection dynamics and their broader implications in an influenza infection and other viral infections [21].

Previously, influenza within-host models often use a mass-action term to describe the infection process by making the assumption of homogenous mixing of cells and viruses [12,16,22]. We show that in the model assuming homogenous mixing (i.e. the HM model), as in previous models, a low-frequency of co-infection and a negligible contribution of SIPs towards total viral load are predicted during the exponential viral growth after infection. However, empirical evidence of influenza’s spatial heterogeneity has been found in humans [42], ferrets [34], and mice [31–33] (see Ref. [23] for a comprehensive overview of influenza’s spatial heterogeneity). These findings showed that foci of infected cells form during an infection, providing a limited range of uninfected cells that newly produced virions can reach. We analyzed a model assuming that viruses can only infect a limited number of target cells (i.e. the TS model) and found that the model predicts a high-frequency of co-infection and a large contribution of SIPs towards viral growth and total viral load, consistent with several previous experimental observations [8,37,38].

Reassortment is common among seasonal IAV strains, potentially contributing to an increased severity of seasonal epidemics [43–45] and to the emergence of pandemic IAV strains [46]. Fonville el al. recently showed that SIPs enhance IAV reassortment, a fundamental process driving IAV evolution and adaptation [11]. The prediction of our model, that co-infection is frequent and SIPs contribute substantially towards the total viral load, thus has important implications in understanding viral genetic diversity, reassortment, adaptation and drug resistance development. Our model here can be a useful tool to provide an upper bound estimate of the extent of coinfection and reassortment in a host. Ignoring SIPs (i.e. the majority of the virus population), however, would lead to an imprecise estimate of the active viral population size, which would in turn lead to an underestimate of the viral diversity in the within-host viral population and the probability of drug resistance (as demonstrated in Perelson et al. [47]), and the frequency of reassortment.

The TS model predicts that the exponential viral growth is mostly driven by virions produced by coinfected cells or multiply-infected cells, due to frequent coinfection (although we do not explicitly track cells infected by more than two virions), while previous models predict that viral growth relies on the virion production rate from singly infected cells. This implies that, although the viral load predictions from the TS model look similar to those of previous models [12], the infection mechanisms and the parameters obtained from fitting the TS model to data are different from what is suggested by previous models. The distinction between infection mechanisms is important when connecting experimental measurements and mathematical models, especially since the virion production rate of a cell may depend on the number of virions that have entered it [26,27]. Thus, our results indicate that experimentally determining the relationship between the viral input (i.e. the multiplicity of infection, MOI) and viral output of an infected cell (the consequences of MOI) and appropriate mathematical models that incorporate these quantities (for example, Ref. [41]) will be crucial to correctly interpret drivers of viral load dynamics. Further, another consequence of frequent coinfection is that the viral growth rate becomes relatively insensitive to changes in the fraction of SIP production. This may explain the substantial variation in SIP production between IAV strains and the counterintuitive observation that a virus that produces more SIPs can actually be more fit than a virus that produces fewer SIPs *in vivo* [39].

Despite the ability to reproduce a wealth of dynamics not seen in previous models, we acknowledge that the TS model has limitations. One such limitation is that our model assumes that cells that are infected with more than two virions are phenotypically identical to those infected with only two virions. This assumption allows us to model cells with any number of co-infecting virions within them while only explicitly modeling the doubly infected cell populations (E_2_ and I_2_). Modeling this system without this assumption would require either a larger, more complicated model [18,19,48] or an alternate model formulation [41].

Our work provides a simple framework to consider the impact of spatial heterogeneity and coinfection on viral dynamics. More broadly, this framework is suitable and can be easily adapted to understand and predict the impacts of reassortment, the origins and consequences of genetic and genomic diversity within viral populations [7], and the process of drug resistance and adaptation for influenza and other viruses [21]. Furthermore, there is a growing interest in developing defective interfering particle (DIP) based therapies recently [49–53]. Given the critical dependence that DIPs have on co-infection to reproduce, our model serves a suitable framework to evaluate the efficacy of DIP-based therapies across a variety of acute viral infections.

## Supporting Figure Captions

**Fig. S1.**
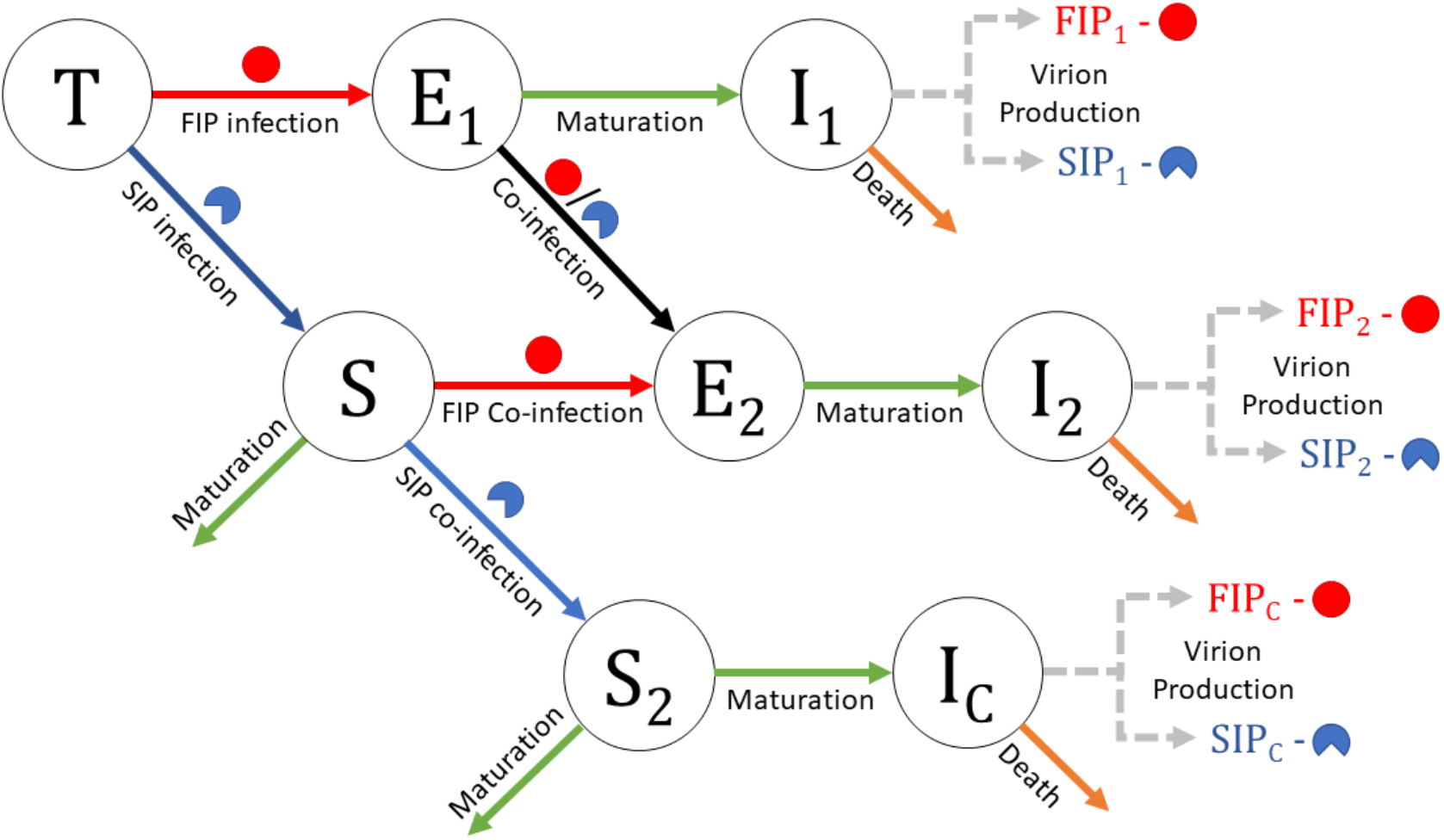
Schematic of expanded model used to track virions created from multiplicity reactivation. *S_2_* cells have been infected with two SIPs. They then mature to *I_C_*, which are virion producing cells derived from complementation (or multiplicity reactivation). In this formulation, *E*_1_ cells are cells that contain both a FIP and a SIP, which mature to *I_2_* cells that produce virions. Thus, we can determine the fraction of virions that are due to complementation by the following equation: 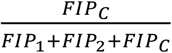. In this formulation, the proportion of co-infected cells is given by 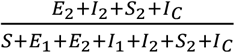 (note that this quantity is the same as 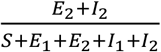 in the original schematic in Fig. 1).

**Fig. S2.**
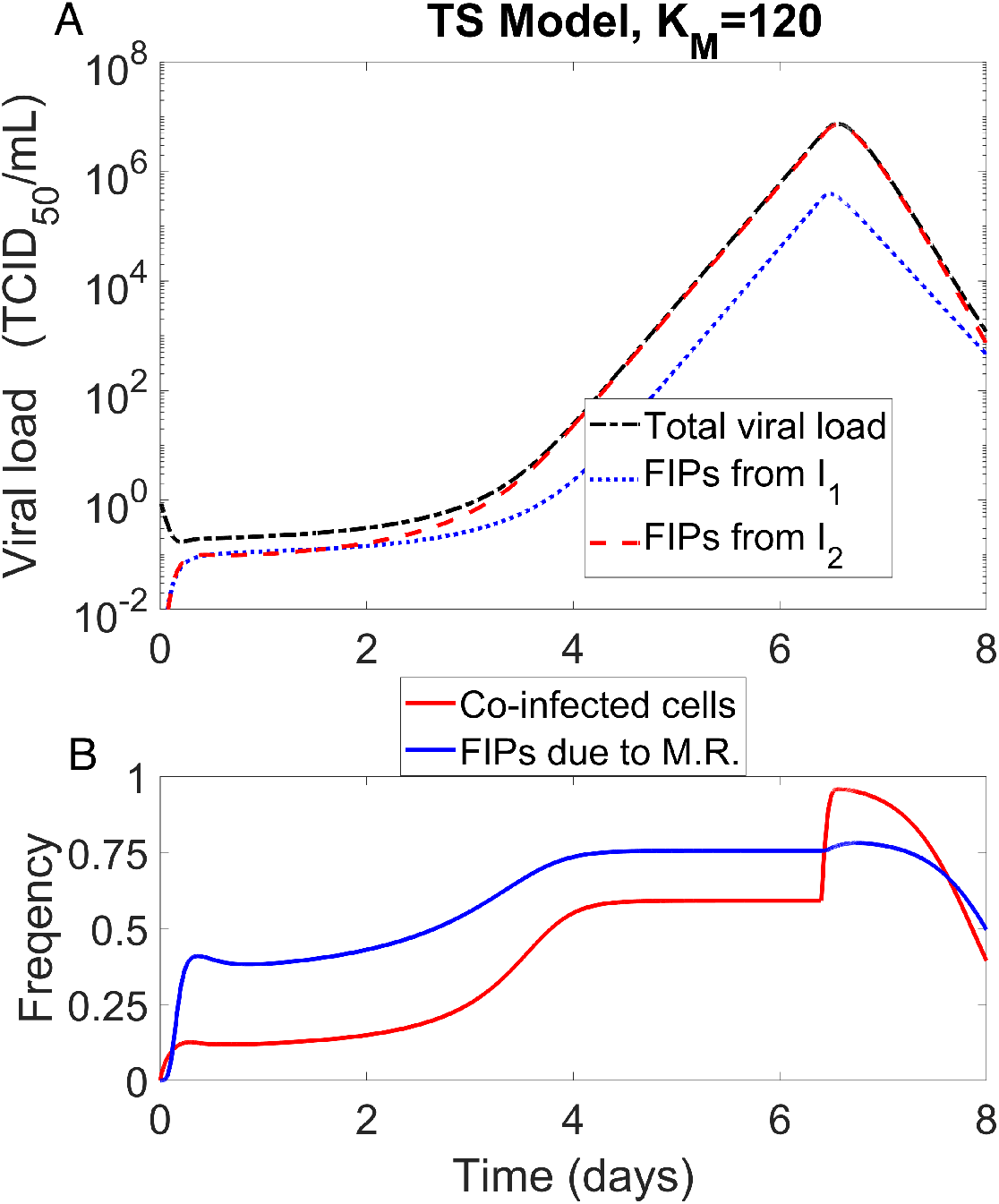
Viral load dynamics and proportion of co-infected cells predicted by the TS model (Eqns. 2) with *K_M_* = 120 cells. Viral load dynamics. **(A)** The total viral load is the concentration of fully infectious virus from all sources. FIPs from *I*_1_ and *I*_2_ (blue and red lines, respectively) are the densities of fully infectious virus produced by singly and doubly infected virion producing cells, respectively. **(B)** The frequency of infected cells that are co-infected (red line) and the frequency of fully infectious virions produced that are due to multiplicity reactivation of SIPs are plotted (blue line; calculations for this quantity are explained in caption of Fig. S1). Multiplicity reactivation occurs when a SIP co-infects a SIP infected cell (*S*), resulting in viral production. The parameters used are from patient 5 in Table 1, except *K_M_* = 120 cells.

**Fig. S3.**
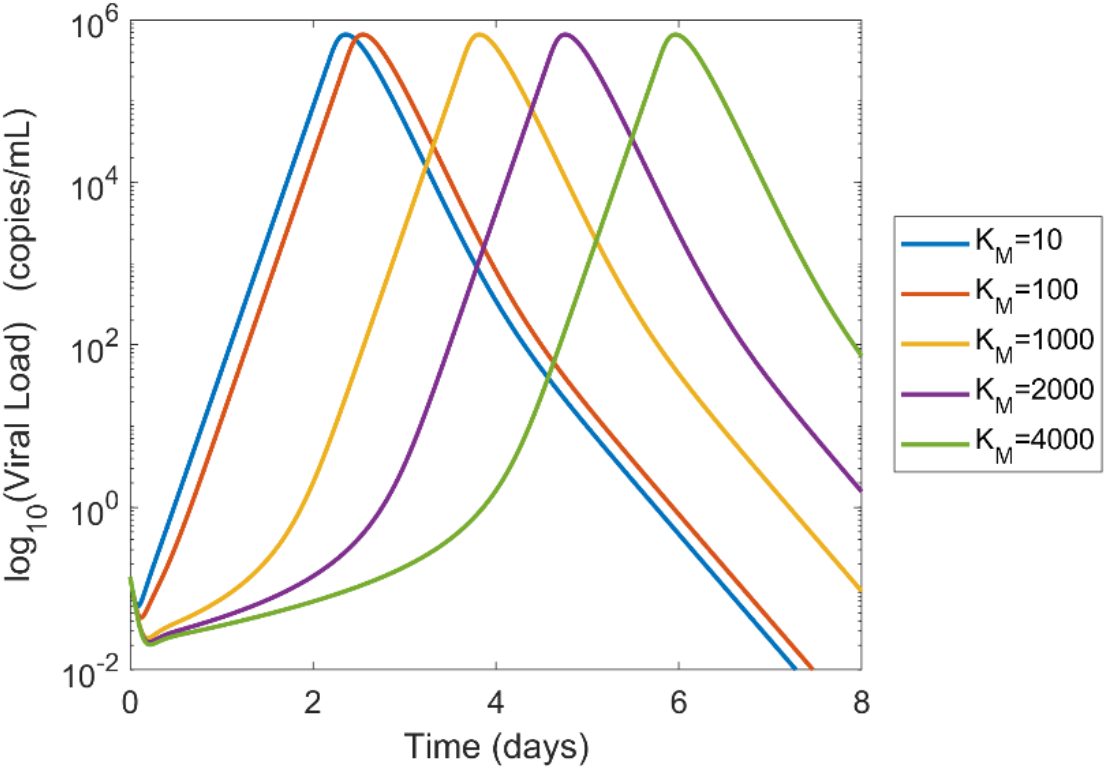
Simulations showing how *K_M_* affects the length of the first phase of growth. We see that increasing the value of *K_M_* increases the length of the first phase of growth. All parameters used (except *K_M_*) are from Patient 5 in Table 1.

